# Lower pathogenicity but similar genital shedding of monkeypox virus clade Ib compared to clade Ia in a rodent model

**DOI:** 10.64898/2025.12.09.693236

**Authors:** Franziska K. Kaiser, Shane Gallogly, Reshma Koolaparambil Mukesh, Missiani Ochwoto, Claude Kwe Yinda, Sarah van Tol, Jonathan Schulz, Kailin Hawes, Brown Bulloch, Anthony McBain, Trenton Bushmaker, Atsushi Okumura, Jessica Prado-Smith, Brian J. Smith, Kyle Rosenke, Greg Saturday, Vincent J. Munster

**Author notes:** Corresponding author: Vincent J. Munster, 903 South 4th Street, Hamilton, MT 59840, USA.

## Abstract

We compared clinical disease, virus shedding and dissemination following genital MPXV clade Ia and Ib infections in *Mastomys natalensis*. MPXV clade Ib resulted in a milder clinical disease but a similar shedding profile to clade Ia. These findings indicate a prolonged pre-symptomatic or prodromal shedding period in clade Ib infections.

## Introduction

In 2024, a new subclade, MPXV clade Ib, emerged in the Democratic Republic of the Congo (DRC)^1^. This is raising concerns, as monkeypox virus (MPXV) clade I has been associated historically with more severe disease^2^.

In addition to clade Ib, several distinct clade Ia lineages of MPXV are circulating in DRC simultaneously^3^. However, only MPXV clade Ib has resulted in a broad international spread, with local circulation in the USA, Europe and Asia^4,5^.

Similar to the 2022 MPXV IIb outbreak, the rapid spread of the new clade Ib has been facilitated by sexual contact and household transmission^6,7^. Yet, limited information is available on the pathogenicity and shedding dynamics of clade Ib MPXV compared to clade Ia. Previously, we demonstrated that infection with MPXV clade IIb via the genital mucosae increases shedding and transmission in the multimammate rat (*Mastomys natalensis*) model^8^.

Here, we compare MPXV clade Ia and Ib clinical disease, pathogenicity and virus shedding kinetics in a vaginal infection model to mimic sexual exposure routes.

## The study

### MPXV clade Ib results in milder clinical disease phenotype compared to clade Ia

To compare the pathogenicity of clade Ia and Ib (MPXV Zaire 79 (V79-I-005) and hMPXV/USA/CA/CDC/2024), we inoculated multimammate rats (*Mastomys natalensis; n=8*) vaginally with (5×10^4^ PFU or 5×10^2^ PFU) of the two MPXV subclades. Four animals per group were euthanized for a scheduled necropsy on day 7, and the remaining animals were monitored for up to 21 days or until they met humane endpoint criteria. Survival rates were 25% for clade Ia high dose infections, 75% for clade Ia low dose and clade Ib high dose infections, and 100% for clade Ib low dose infections (Figure 1A). Animals of the clade Ia high dose group were the only animals with a strong temperature increase, peaking on day 10 (Figure 1B). Relevant weight loss was only detected following high dose clade Ia infection (Figure 1C-F).

**Figure 1.**
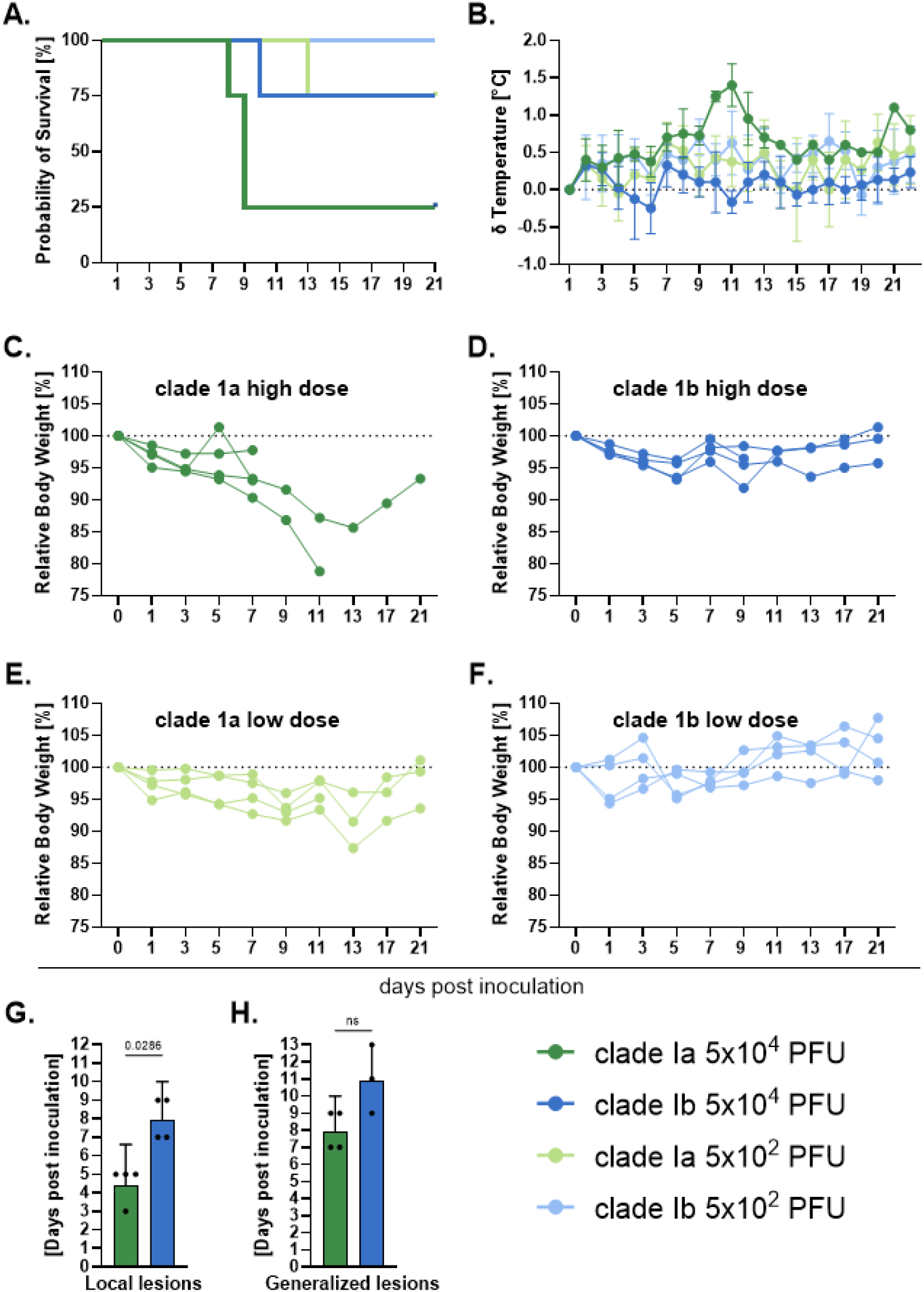
Higher, dose-dependent disease severity with MPXV clade Ia compared to clade Ib. Four animals per group were vaginally inoculated with 5×10^4^ (high dose) or 5×10^2^ (low dose) plaque forming units (PFU) MPXV clade Ia (MPXV Zaire 79, V79-I-005) or clade Ib (hMPXV/USA/CA/CDC/2024). Clinical exams were performed, and body weight was recorded at 1, 3, 5, 7, 9, 11, 13, 17, and 21 days post inoculation (dpi). **A**.Kaplan-Meier survival curve shows the probability of survival for each subclade and dose. **B**. Temperature changes were measured by implanted transponders at 1.30am during the animals’ noctuid activity phase; symbols and error bars display mean and standard deviation. **C-F**. Relative body weight changes compared to the day of inoculation, following clade Ia high dose (C), clade Ia low dose (D), clade Ib high dose (E), and clade Ib low dose (F) infections. **G**. Days until first signs of infection were visible at the inoculation site following high dose inoculation (*n=4*). These lesions included redness, swelling, and vaginal discharge. Statistical significance was calculated by Mann-Whitney U test. **H**. Days until first generalized lesions were visible in high dose groups. These included skin lesions around the base of the tail, at the eyelids or disseminated skin lesions (*n=4* for clade Ia, *n=3* for clade Ib). Statistical significance was calculated by Mann-Whitney U test.

### Comparable clade Ia and Ib infectious MPXV shedding at the site of inoculation

Next, we compared genital, oropharyngeal, nasal, and rectal virus shedding following vaginal inoculation with clade Ia or Ib, (5×10^4^ PFU or 5×10^2^ PFU). Shedding of infectious virus at the inoculation site did not differ depending on the subclade (Figure 2). At the non-inoculation sites (oropharyngeal, nasal, and rectal), overall shedding as calculated by the area under the curve (AUC) analysis showed a trend to be higher following high dose clade Ia inoculation, compared to clade Ib.

**Figure 2.**
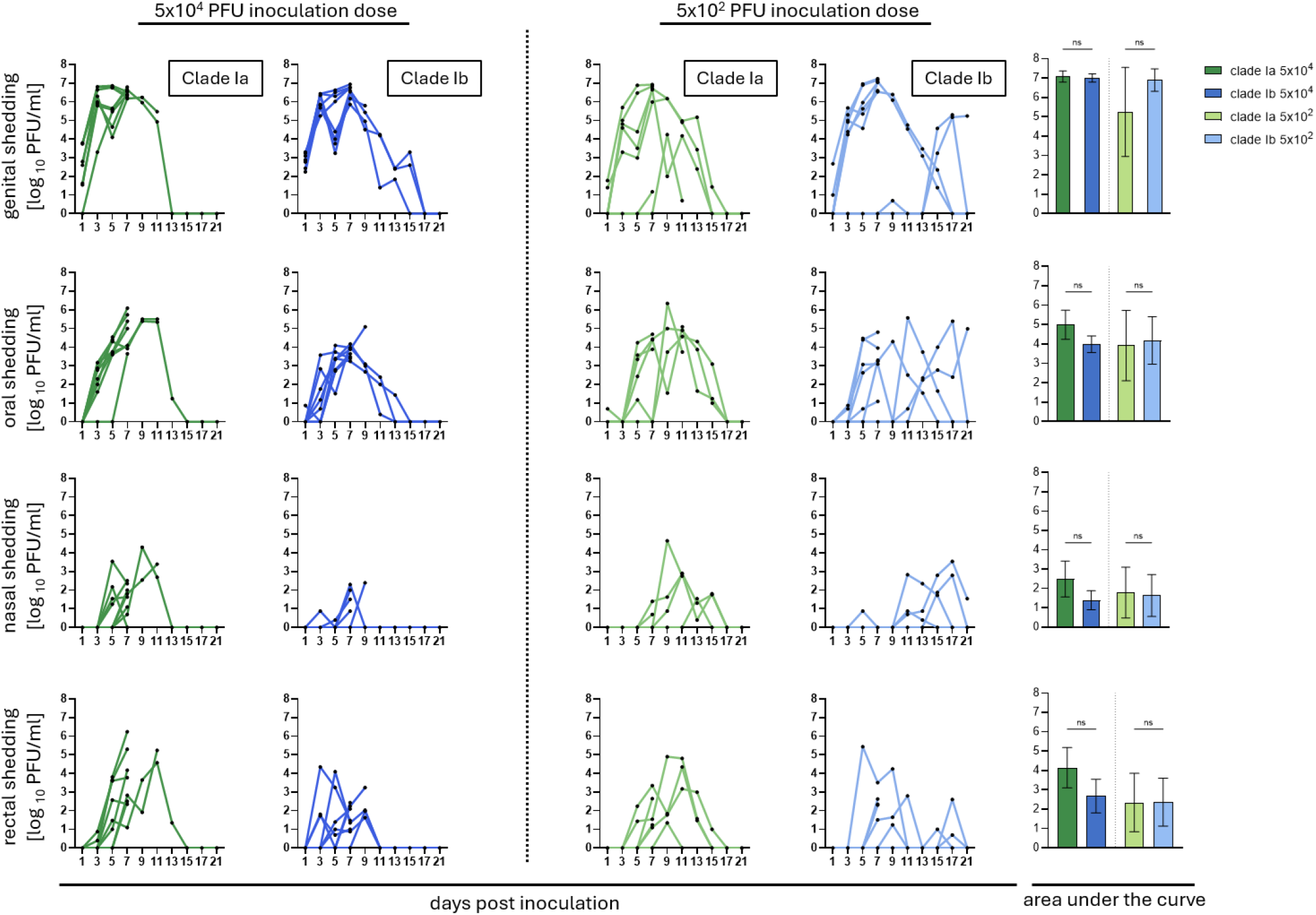
Comparable infectious MPXV shedding kinetics following MPXV clade Ia or Ib at inoculation site. Vaginal, oropharyngeal, nasal, and rectal swab samples were collected on day 1, 3, 5, 7, 9, 11, 13, 15, 17, and 21 post vaginal inoculation or until they met endpoint criteria. Infectious virus particles [PFU/ml] were quantified by plaque assay. Lines represent individual animals. Areas under the curve (AUCs) were calculated for each shedding route (geometric mean; 95% CI). AUC results were tested for statistical significance using a Kruskal-Wallis test.

Vaginal shedding was detected on 1 day post-inoculation (dpi) in all but one animal among both subclades in the high dose inoculation groups with 2.3×10^3^ (SD 2.7×10^3^) PFU/ml in clade Ia and 9.4×10^2^ (SD 6.2×10^2^) PFU/ml in clade Ib (mean 1 dpi). Oropharyngeal, nasal, and rectal shedding were first observed between 3 and 5 dpi for most animals.

Overall, a lower inoculation dose did not result in reduced virus shedding. However, the infection efficiency was lower, and some animals in the low dose inoculation groups did not start shedding until 7 dpi. These animals were co-housed with others that did shed infectious virus from 1 dpi onwards. Low dose inoculations caused a 62.5% (5/8) infection efficiency for clade Ia and 75% (6/8) infection efficiency for clade Ib, respectively (Supplemental Figure 1). This initial infection efficiency was calculated as the percentage of animals productively infected with MPXV at 5 dpi. 5×10^4^ PFU inoculation of clade Ia or Ib resulted in a 100% (8/8 per subclade) infection efficiency.

### Virus dissemination and systemic spread following clade Ia and clade Ib infections

We compared virus dissemination in different organs (skin, lung, liver, spleen, kidney, bladder, reproductive tissue) at 7dpi in four animals per group. High dose inoculation of MPXV clade Ia and Ib resulted in systemic virus spread and detection in all tested organs (Figure 3A). Reproductive tissue and skin tissue had the highest quantity of MPXV genomic material. Reproductive tissue reached 1.1×10^11^ (SD 1×10^11^) for clade Ia and 3.4×10^7^ (SD 3.9×10^7^) copies/gram for clade Ib, and skin tissue 4.6×10^9^ (SD 7×10^9^) for clade Ia and 9.9×10^7^ (SD 7.5×10^7^) copies/gram for clade Ib. When comparing overall virus loads in all tested organs, clade Ia high dose resulted in an average of 1.6×10^10^ (SD 5.3×10^10^) copies/gram and clade Ib in 2.9×10^7^ (SD 4.5×10^7^) copies/gram. Variability in virus dissemination on 7 dpi was higher in the low dose inoculation groups (Figure 3B).

**Figure 3.**
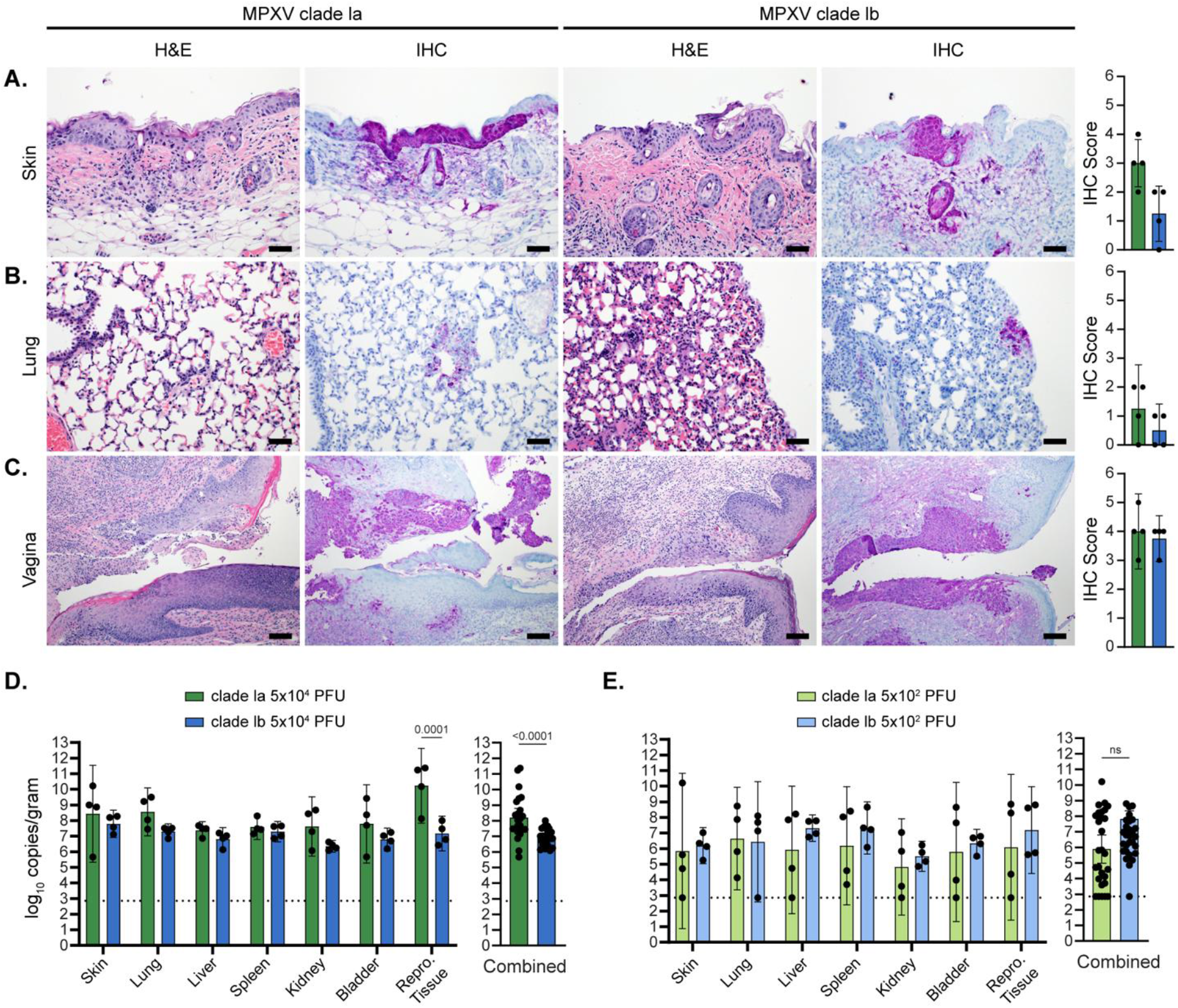
Higher virus loads in systemic organs on day 7 following MPXV clade Ia infection. 4 animals per group were vaginally inoculated with PFU mpox clade Ia or clade Ib. Panel **A.-C**. are representative hematoxylin eosin-stained (H&E) or immunohistochemistry (IHC) images of skin, lung, or vaginal mucosa. Graphs display semiquantitative histological score based on IHC antigen presence. **A**. Skin clade Ia displayed multifocal, moderate subacute dermatitis with moderate immunoreactivity of epithelial cells, fibroblasts and inflammatory cells while Ib has mild inflammation and scattered immunoreactivity. **B**. Lung clade Ia & Ib had one animal in each group with subacute pulmonary infiltrates and scattered immunoreactivity of pneumocytes. **C**. Vaginal inoculation site clade Ia & Ib displayed marked to severe subacute vaginitis with abundant immunoreactivity of epithelial cells, fibroblasts and inflammatory cells in each group. **A/B** (200x magnification) Scale bars equal 50µm; **C** (100x magnification) Scale bars equal 100µm. Panel **D**. displays copy numbers in different organs following 5×10^4^ PFU (high dose) and **E**. 5×10^2^ PFU (low dose) inoculation dose. The bars depict the geometric means, error bars show 95% confidence interval, and circles represent individual animals. Statistical significance was tested via two-way ANOVA followed by Sidak’s multiple comparison test for organ tissues and Mann-Whitney U test for combined values.

### Presymptomatic and prodromic shedding of infectious MPXV

Next, we compared the temporal occurrence of infection-associated parameters after high dose (5×10^4^ PFU) vaginal inoculation during the 21-day observation period (Figure 4). We observed early onset of high MPXV shedding on 1 dpi and with a plateau phase between 3 and 7 dpi, identical for clade Ia and Ib. Localized signs of inflammation appeared 3 dpi in 25% (1/4) of animals for clade Ia and by 5 dpi all animals displayed signs of localized inflammation (Figure 4A). In contrast, after Ib infection localized signs of inflammation started 7 dpi for 50% (2/4) of the clade Ib infected animals and by 9 dpi all animals displayed signs of localized inflammation (Figure 4B). Generalized skin lesions appeared in 50% (2/4) of animals in clade Ia group on 7 dpi and in all (4/4) animals between 9-13 dpi. Clade Ib infected animals displayed generalized lesions in 25% (1/4) animals on day 9, 33.3% (1/3) on day 11 and 66.7% (2/3) on day 13. The temperature increase peaked on 9 dpi following clade Ia infection, no fever peak was observed in the clade Ib group. Similarly, the body weight loss peaked on 11 dpi for clade Ia and did not present a clear trend in clade Ib.

**Figure 4.**
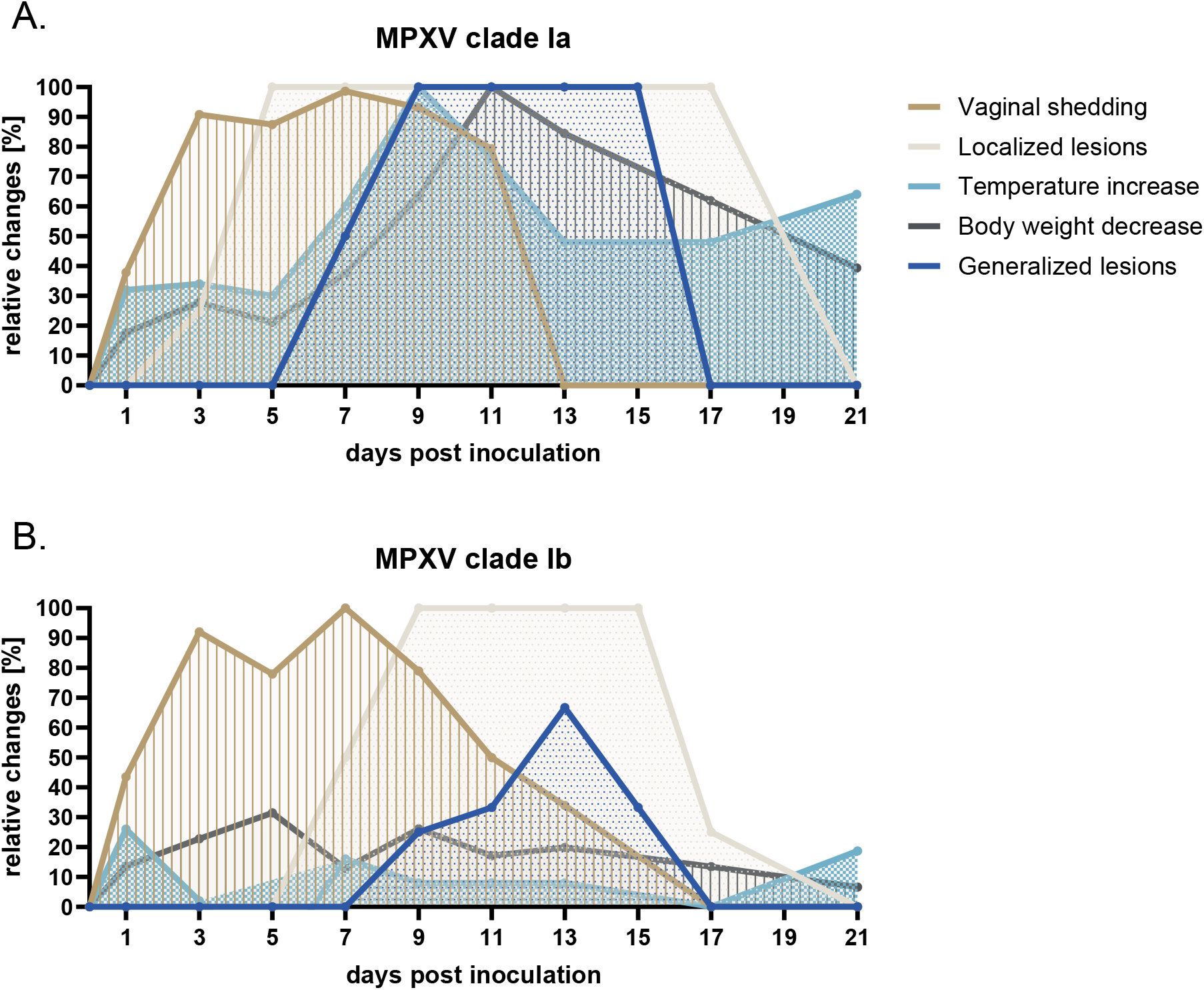
Prolonged presymptomatic virus shedding for MPXV clade Ib compared to Ia. The graphs display decrease in body weight, increase in body temperature, infectious vaginal shedding, percentage of animals with visible signs of localized inflammation and generalized skin lesions following vaginal inoculation with 5×10^4^ PFU. **A**. Correlation for clade Ia. **B**. Correlation for clade Ib. For each parameter the mean was calculated per timepoint from all animals per group. The peak in change compared to start value was equated to 100%. Infectious shedding data was log-transformed prior to calculations. The clade Ia group consists of 4 animals until day 7 (*n=8* for shedding data until day 7), 2 animals until day 11 and 1 animal until day 21 (due to reaching humane endpoint criteria). The clade Ib group includes 4 animals until day 7 (*n=8* for shedding data until day 7), 4 animals until day 9 and 3 animals until day 21.

## Discussion

This study investigated the clinical disease, pathogenicity, and virus shedding profile of the newly emerged MPXV clade Ib in comparison to clade Ia in the *M. natalensis* model. Vaginal infections with clade Ib resulted in a milder clinical disease and higher survival rate compared to clade Ia in this study. Additionally, virus dissemination in organ tissues and shedding of infectious virus from non-inoculation sites were showed a trend to be reduced following clade Ib versus clade Ia infection. These results are comparable with human data, which indicate a milder disease phenotype and less systemic complications with clade Ib ^9-13^.

Compared to a study with MPXV clade IIb in *M. natalensis*, clade IIb MPXV was less pathogenic than clade Ib (and Ia) but showed a similarly high susceptibility of the anogenital mucosae. This suggests that anogenital mucosal surfaces are highly susceptible for initial MPXV infection, irrespective of the clade^8^.

To determine the potential for asymptomatic transmission during the incubation period, we analyzed the temporal relationship of clinical signs with MPXV shedding. Clade Ia and Ib displayed comparable amounts of virus shedding from vaginal mucosae. However, the onset of clinical signs was significantly slower with clade Ib. Prolonged pre-symptomatic or prodromal virus shedding, combined with milder disease, could be contributing to the epidemiological dynamics of clade Ib emergence.

In conclusion, MPXV clade Ib displayed milder clinical disease phenotype compared to clade Ia in a rodent model. Clade Ib infection displayed a prolonged period of pre-symptomatic and prodromal virus shedding.

## Data and materials availability

All data supporting the findings of this study have been deposited on Figshare and will be made publicly available upon acceptance. A DOI will be provided in the final published version.

## Acknowledgements

We thank the Poxvirus and Rabies Branch at the Centers for Disease Control and Prevention (CDC), Georgia, USA, for helpful discussion and providing a MPXV isolate; BEI, NIAID resources for supplying MPXV isolates; the Office of the Chief, RML, NIAID, NIH for their support within the high containment facility; the animal care staff and veterinarians of the Rocky Mountain Veterinary Branch, NIAID, NIH for their assistance during the study; and the Visual and Medical Arts Section, RML, NIAID, NIH for their help in preparing the graphical illustration.

## Funding Statement

This research was supported by the Intramural Research Program of the National Institutes of Health (NIH). The contributions of the NIH authors are considered Works of the United States Government. The findings and conclusions presented in this paper are those of the authors and do not necessarily reflect the views of the NIH or the U.S. Department of Health and Human Services.

## Conflict of Interest

The authors declare no conflict of interest.

## Notes

### Competing Interest Statement

The authors have declared no competing interest.

