## Appendix for "Lower pathogenicity but similar genital shedding of monkeypox virus clade Ib compared to clade Ia in a rodent model"

### Materials and Methods

#### Ethics statement and biosafety.

Animal experiments were performed after obtaining approval from the Rocky Mountain Laboratories (RML) Institutional Animal Care and Use Committee. The animal holding facility was accredited by the Association for Assessment and Accreditation of Laboratory Animal Care with climate-controlled animal holding rooms with fixed 12-hour light-dark cycles. Experiments were conducted following the Guide for the Care and Use of Laboratory Animals, the Animal Welfare Act, US Department of Agriculture, and the US Public Health Service Policy on Humane Care and Use of Laboratory Animals.

The NIH RML Institutional Biosafety Committee (IBC) approved these experiments and all standard operating and inactivation procedures to be conducted under BSL 3 conditions. This study did not classify as Dual Use Research of Concern (DURC) or Enhanced Potential Pandemic Pathogens (ePPP) project according to the NIH Dual Use Research of Concern-Institutional Review Entity (DURC-IRE).

#### Viruses and cells.

Vero E6 cells were provided by R. Baric (also available as VERO C1008 from ATCC (CRL-1586, https://www.atcc.org/products/all/crl-1586.aspx)) and cytochrome b verified. Virus propagation and quantification of infectious virus by plaque assay were conducted on vero cells in Dulbecco's Modified Eagle Medium (DMEM) supplemented with 2% fetal bovine serum (FBS), 1 mM L-glutamine, 50 U $ml^{-1}$ penicillin, and 50 µg $ml^{-1}$ streptomycin (DMEM2). Prior to the experiments, Vero E6 cells were maintained in DMEM supplemented with 10% FBS, 1 mM L-glutamine, 50 U $ml^{-1}$ penicillin, and 50 µg $ml^{-1}$ streptomycin. Mpox clade Ib isolate hMPXV/USA/CA/CDC/2024 was provided by the CDC. The mpox clade Ia isolate MPXV Zaire 79 (V79-I-005) was obtained from BEI resources (https://www.beiresources.org/Catalog/animalViruses/NR-2324.aspx). Both viruses were sequenced prior to this study, to exclude contaminations and confirm the sequence.

**Animal study.**

*Mastomys natalensis* (10-20 weeks old) were obtained from the RML in-house breeding colony[1]. During the study, animals were group-housed in ventilated disposable cages (Innocage IVC Rat Caging System, Innovive) and provided with ad libitum rodent feed and water. All experimental procedures were done on anesthetized animals under isoflurane inhalation anesthesia.

Prior to study start, we calculated group sizes, using a t-test (alpha = 0.5, power = 0.8, website https://homepage.univie.ac.at/robin.ristl/samplesize.php?test=ttest) using estimated outcome virus titers. Animals were then assigned to four experimental groups of *n=8* (all females) and were allowed to adapt to husbandry conditions in the high-containment environment for 10 days prior to inoculation. At the time point of transferal into the high-containment animal facility, *M. natalensis* were subcutaneously implanted with transponders to monitor body temperatures (UCT-2112, Unified Information Devices (UID)).

On day 0, the four experimental groups were vaginally inoculated with $5x10^4$ PFU or $5x10^2$ PFU MPXV clade Ia or Ib, respectively. Anesthetized animals were positioned on their backs while a dry sampling swab (Polyester tipped applicator, 25-800 1PD, Puritan, ME, USA) was used to remove mucus from their vaginas and inflict microabrasions on the vaginal mucosa by gently rotating the swab[2]. The virus suspension was then carefully pipetted into the vagina in 40µl sterile cell culture medium (DMEM2). Animals were then monitored at least once daily for clinical signs of illness, including but not limited to reduced general condition, segregation from their cage mates, reduced food and water intake, unkempt fur, and hunched posture. Full clinical exams, swab collection, and weighing were done on day 1, 3, 5, 7, 9, 11, 13, 15, 17, 19, and 21. *N=4* animals per group were taken for a scheduled necropsy on day 7 post inoculation and the other four animals at the study endpoint on day 21.

The monitoring during exams had a special emphasis on signs of localized inflammation at the inoculation site and on the occurrence of generalized skin lesions. Localized inflammation mostly presented with redness, swelling, and vaginal discharge. Generalized skin lesions occurred predominantly as rashes and pustules on the base of the tail, eyelids, or widely disseminated pox lesions.

**MPXV quantitative real-time PCR.**

RNA was extracted from animal tissues using a QIAamp Viral RNA Kit (Qiagen) according to the manufacturer's description. Organ tissue was weighed and then homogenized in RLT buffer. MPXV nucleic acids were quantified by using a MPXV-specific qRT-PCR assay for the detection of G2R gene coding sequence[3]. The forward primer 5'-GGAAAATGTAAAGACAACGAATACAG-3', reverse primer 5'-GCTATCACATAATCTGGAAGCGTA-3' and probe 5'-FAM-AAGCCGTAATCTATGTTGTCTATCGTGTCC-3'-BHQ1 were used as reaction mix in a TaqMan Fast Virus One-Step Mastermix (Applied Biosystems) with 15µl reaction mix and 5µl extracted nucleic acids. For nucleic acid quantification, we used a QuantStudio 3 Flex Real-Time PCR System (Applied Biosystems) according to instructions of the manufacturer. Serial dilutions of MPXV standards with known copy numbers were included in each PCR run to generate a standard curve and calculate copy numbers.

**Infectious virus quantification by endpoint plaque assay.**

Swab samples were inserted into 1ml DMEM2 following their collection and stored at -80°C for downstream processing. Ten-fold serial dilutions in DMEM2 were added to 48-well plates coated with Vero E6 cells. Cell plates were incubated for four days at 37°C, 5% $CO_2$ until fixated with 10% formaldehyde for 10min. Plates were stained with 1% crystal violet solution for 10min, followed by rinsing and drying for 24h. After drying, the plaques were counted and plaque forming units per ml (PFU/ml) were calculated.

### Histopathology

Tissues were placed in cassettes and fixed for a minimum of 7 days in 10% neutral buffered formalin. Downstream processing was done with a Sakura VIP-6 Tissue Tek on a 12-hour automated schedule using a graded series of ethanol, xylene, and PureAffin. Section of 5 µm were cut of embedded tissue and dried overnight at 42°C, prior to processing. MPXV antigen presence was detected using GeneTex Vaccinia virus antibody (#GTX36578) at a dilution of 1:2000. Vector Laboratories ImmPRESS VR horse anti-rabbit IgG polymer (# MP-6401) was used as a secondary antibody. For negative controls, replicate sections from each block were deparaffinized and stained in parallel following an identical protocol, with the primary antibody replaced by Vector Laboratories rabbit IgG (#I-1000-5) at a dilution of 1:2500. For tissue staining, the Discovery Ultra automated stainer (Ventana Medical Systems) was utilized with a Roche Tissue Diagnostics Discovery purple kit (#760-229). A board-certified, blinded pathologist performed histopathological assessment.

### Statistical analysis

Data analysis and test for statistical significance were performed using GraphPad Prism software version 10.2.0. Statistical tests and p-values are indicated where appropriate. *p*-values>0.05 were interpretated as not significant (*ns*).

**Supplemental Figures**

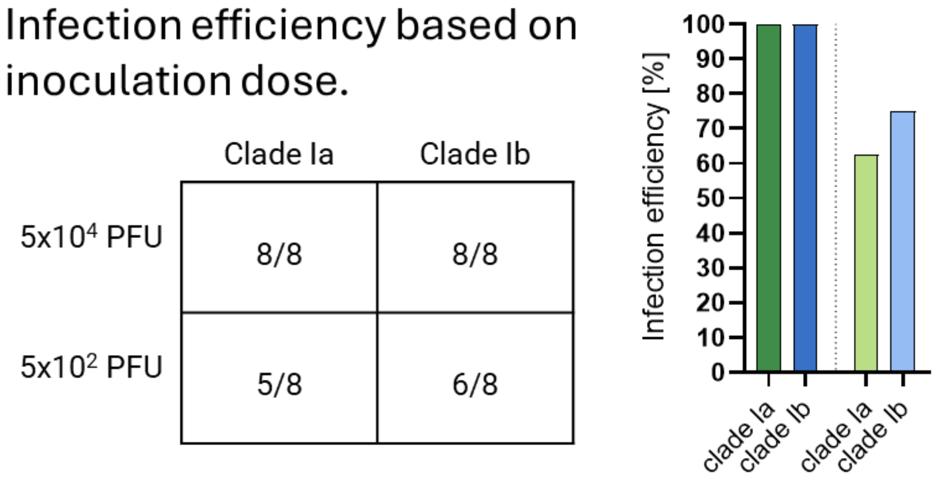


**Supplemental Figure 1: Initial infection efficiency following vaginal inoculation**. Infection doses are $5x10^4$ PFU (high dose) and $5x10^2$ PFU (low dose) mpox virus subclade Ia or Ib. The infection rate represents the percentage of animals with detectable vaginal virus shedding on day 5 (n=8 per group). *n=8* animals were included per virus clade and inoculation dose. Choosing the data from day 5 ensures the initial infection rate was captured before animals acquired the infection from co-housed cage mates.

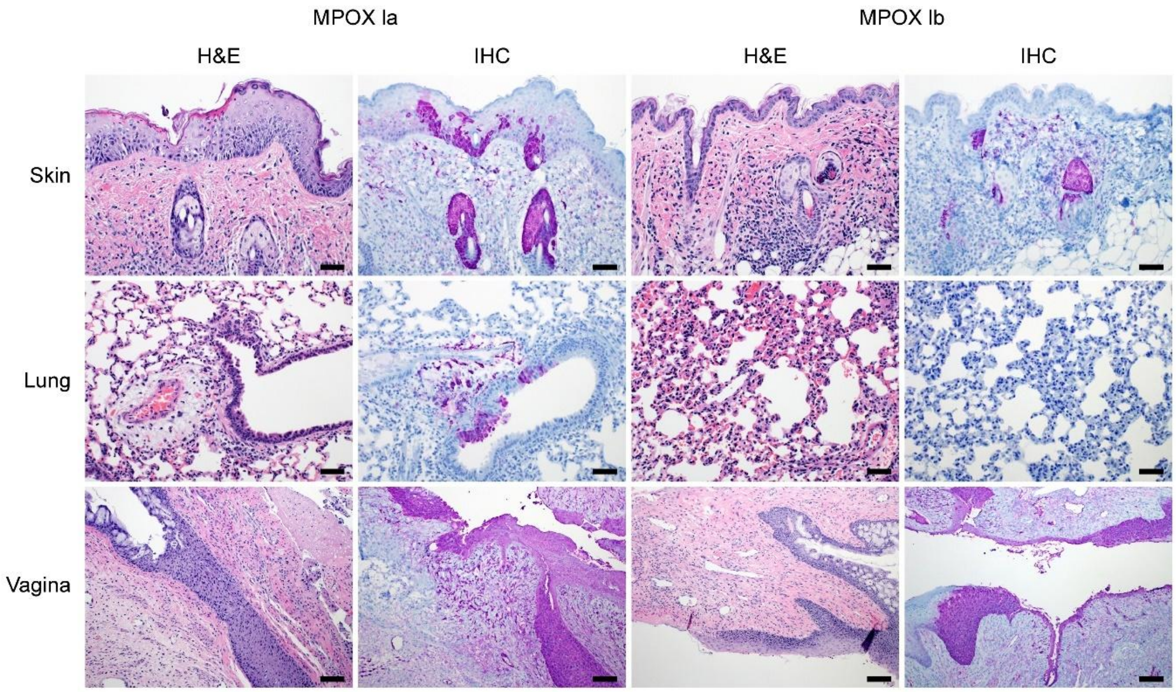


**Supplemental Figure 2: Histopathological findings in low dose MPXV infections**

4 animals per group were vaginally inoculated with PFU mpox clade Ia or clade Ib. Panel **A.-C.** are representative hematoxylin eosin-stained (H&E) or immunohistochemistry (IHC) images of skin, lung, or vaginal mucosa. Graphs display semiquantitative histological score based on IHC antigen presence. **A**. Skin clade Ia two animals displayed mild subacute dermatitis with scattered immunoreactivity of epithelial cells, fibroblasts and inflammatory cells while Ib has one animal with mild inflammation and rare immunoreactivity. **B.** Lung clade Ia had one animal with subacute pulmonary infiltrates and scattered immunoreactivity of pneumocytes and bronchiolar epithelium. Clade Ib did not have pulmonary lesions or immunoreactivity **C.** Vaginal inoculation site clade Ia & Ib displayed moderate to marked subacute vaginitis with abundant immunoreactivity of epithelial cells, fibroblasts and inflammatory cells in each group. **A/B** (200x magnification) Scale bars equal 50μm; **C** (100x magnification) Scale bars equal 100μm.
